# Mechanoaccumulation of non-muscle myosin IIB during mitosis requires its translocation activity

**DOI:** 10.1101/2021.09.08.459548

**Authors:** Chao Wang, Jingjing Ding, Qiaodong Wei, Shoukang Du, Xiaobo Gong, Ting Gang Chew

**Author notes:** Correspondence to: Ting Gang Chew. Co-first authors (Chao Wang, Jingjing Ding).

## Abstract

Non-muscle myosin II (NMII) is a force-generating mechanosensitive enzyme that responds to mechanical forces exerted on cells. Mechanoresponse of NMIIs confers mechanical adaptability to cells growing and dividing in a physically complex microenvironment. In response to mechanical forces, NMIIs mechanoaccumulate at the cell cortex with applied stress. Much less is known about how NMII mechanoaccumulation is mechanistically regulated. In this study, we subject cells in mitosis to compressive forces and show that mitotic cells promote active RhoA mechanoaccumulation, and via ROCK signaling, activate and stabilize NMIIB at the cell cortex. In line with RhoA in activating the myosin motor activity, we further show that the motor activity driving actin filament translocation, but not just the actin-binding function of NMIIB plays a dominant regulatory role in NMIIB mechanoaccumulation. Thus, the motor activity coordinates structural movement and nucleotide state changes to fine-tune actin-binding affinity optimal for NMIIs to generate and respond to forces.

## Introduction

Non-muscle myosin IIs (NMIIs) are a family of actin-based molecular motors that function with actin filaments to generate forces to drive cell morphogenesis, cell migration and cell division (Murrell et al., 2015; Vicente-Manzanares et al., 2009). NMIIs and actin filaments form a layer of contractile cytoskeletal network at the cell cortex underlying the cell membrane to drive changes of cell shape (Kelkar et al., 2020). One key role of NMII’s contractility is to promote mitotic cell rounding in cells transiting from interphase into mitosis, and to sustain a round cell shape throughout mitosis (Nishimura et al., 2019; Ramanathan et al., 2015; Ramkumar and Baum, 2016). Cells that are not able to maintain a round cell shape when dividing in a physically confined environment have defects in their mitotic spindle stability and form multipolar spindles that lead to chromosome mis-segregation and aneuploidy (Cadart et al., 2014; Lancaster et al., 2013; Matthews et al., 2020).

As cells enter mitosis, activation of cyclin-dependent kinase 1 (CDK1) triggers the cytoplasmic shuttling of a Rho guanine nucleotide exchange factor (RhoGEF) Ect2 to activate RhoA, which in turn binds to the cell membrane and recruits its effectors such as anillin, formin and Rho-associated kinase (ROCK) to activate NMII sliding on actin filaments homeostatically maintained at the cell cortex (Chew et al., 2017; Maddox and Burridge, 2003; Matthews et al., 2012; Ramanathan et al., 2015; Rosa et al., 2015). Activation of this signaling cascade results in generation of higher cell surface tension, which leads to mitotic cell rounding crucial in providing an ideal cell geometry to undergo mitosis and cytokinesis (Cattin et al., 2015; Lancaster et al., 2013; Stewart et al., 2011).

NMII consists of two heavy chains, two regulatory light chains (MRLC) and two essential light chains (ELC). The myosin heavy chain has a globular head, a flexible neck structure and an elongated tail region (Sweeney and Houdusse, 2010). NMII is activated when the neck structure is bound by MRLC phosphorylated via RhoA-mediated ROCK signaling (Amano et al., 1996; Garrido-Casado et al., 2021). Further phosphoregulation of the myosin tail region assembles active NMIIs into multimeric myofilaments that have high affinity to actin filaments (Garrido-Casado et al., 2021; Juanes-Garcia et al., 2015). The motor activity in the globular head of myofilaments catalyses nucleotide hydrolysis to form the cross-bridge and to translocate actin filaments via lever arm movement (Sweeney and Houdusse, 2010). Apart from force generation, NMIIs sense and respond to mechanical inputs in cells. In response to mechanical forces, NMIIs accumulate at the cell cortex of the stressed region in cells (Luo et al., 2013). The three mammalian NMII paralogs (NMIIA, NMIIB, NMIIC) accumulate at the region that is mechanically stressed (mechanoaccumulation), with NMIIB showing differential mechanoresponse across cell types and cell cycle phases (Schiffhauer et al., 2016). Mechanoaccumulation of NMIIs is influenced by the length of its neck structure in which an extended length of the neck structure facilitates its accumulation (Luo et al., 2013). Furthermore, phosphoregulation of NMII’s tail region that modulates myofilament assembly states contributes to the ability of NMIIs to accumulate at the cell cortex in response to mechanical stress (Schiffhauer et al., 2019).

Much has been done to understand the force generation mechanism of NMIIs, less is known about how NMII’s mechanoresponse is regulated. In this study, we aimed to understand the mechanism for NMIIB mechanoaccumulation in mitotic cells responding to compressive forces. By using a cell contractility reporter cell line, we found that cells respond to compressive stress during mitosis by accumulating active RhoA at the cell cortex, which in turn leads to higher activation and mechanoaccumulation of NMIIB. Interestingly, further genetic dissection of NMIIB head domain by using mutations that uncouple actin-binding function and motor activity to translocate actin filaments showed that the translocation activity of NMIIB plays a primary role for its ability to undergo mechanoaccumulation. We propose that the NMIIB translocation activity coordinates structural changes to nucleotide state changes to prepare NMIIBs of differential actin-binding affinity, so to achieve mechanoaccumulation at the cell cortex in response to compressive forces during mitosis.

## Results

### Construction and validation of a cell contractility reporter cell line

To study how NMII mechanoresponse is regulated during mitosis, we made a human MCF10A mammary epithelial cell stably expressing fluorescently tagged non-muscle myosin IIB (mEmerald-NMIIB) and active RhoA reporter consisting of the RhoA-binding domain of anillin (mRuby2-AHPH) (Acharya et al., 2018; Beaudet et al., 2020; Cavanaugh et al., 2022; Piekny and Glotzer, 2008; Priya et al., 2015). NMIIB localizes to the cell cortex in the early mitosis and functions downstream of active RhoA to generate cortical tension (Maddox and Burridge, 2003; Taneja et al., 2020). Live-cell microscopy imaging revealed that NMIIB and active RhoA localized to the cell cortex starting from early mitosis and enriched at the division site during mitosis and cytokinesis, consistent with previous studies and their roles in tension generation at the cell cortex during cell division (Fig. 1 A) (Maddox and Burridge, 2003; Piekny and Glotzer, 2008; Taneja et al., 2020). To examine the activity of mEmerald-NMIIB, we treated mitotic cells with low, medium and high concentrations of the low cytotoxic and photostable NMII inhibitor para-nitro-blebbistatin (p-nitroblebb), which lowers the actin affinity of NMII heads (Kepiro et al., 2014; Rauscher et al., 2018). Mitotic cells treated with a low concentration of p-nitroblebb showed punctate cortical NMIIB patterns (Fig. 1 B, insets). Higher concentrations of p-nitroblebb resulted in a decreasing NMIIB intensity at the mitotic cell cortex (Figs. 1 B and C). Analyses of the NMIIB turnover at the mitotic cell cortex with the fluorescence recovery after photobleaching technique (FRAP) showed an increase of the immobile fractions of NMIIB upon increased concentrations of p-nitroblebb (Figs.1 D and E; DMSO: 0.27 ± 0.08; [Low]: 0.45 ± 0.01; [Medium]: 0.50 ± 0.01; [High]: 0.59 ± 0.03). The recovery half time of cortical NMIIB decreased when increasing concentrations of p-nitroblebb were used (Fig. 1 E; DMSO: 20.33 ± 2.09 s; [Low]: 12.61 ± 1.46 s; [Medium]: 11.30 ± 0.59 s; [High]: 9.49 ± 0.05 s). The FRAP analysis suggested that a decrease of NMIIB recovery half time with higher immobile fractions characterizes inhibition of mEmerald-NMIIB function at the cell cortex by the NMII inhibitor.

**Figure 1.**
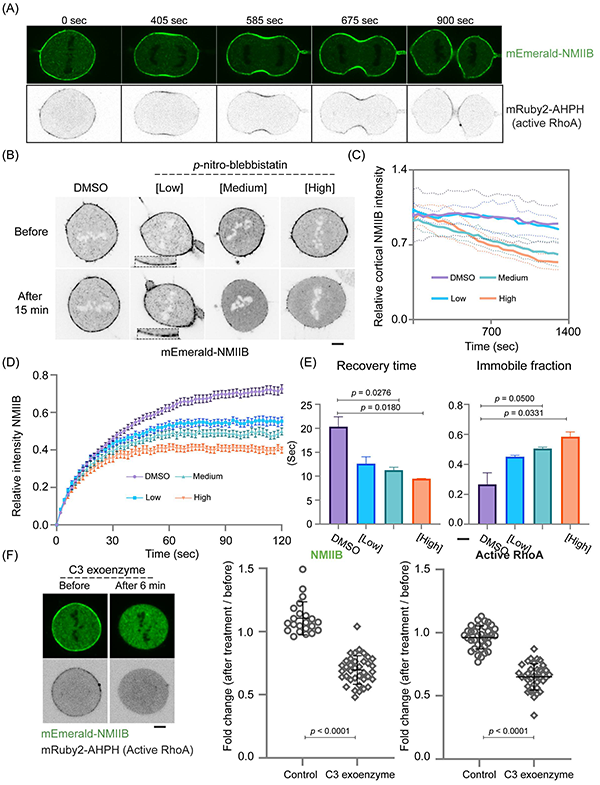
Establishment of a MCF10A cell line stably expressing mEmerald-NMIIB and mRuby2-AHPH (active RhoA). (A) Micrographs of a representative cell showing mEmerald-NMIIB and active RhoA localization during mitosis and cytokinesis. (B) Micrographs of the cortical mEmerald-NMIIB in mitotic cells treated with different concentrations of p-nitroblebb. Cells before and 15 minutes after the treatments were shown. [Low] is 37 μM; [Medium] is 74 μM; [High] is 110 μM. Insets show a zoom-in view of the punctate NMIIB. (C) Relative changes of cortical NMIIB intensity as a function of time. The fluorescence intensity at each time point was normalized to that of in cells before treatments. Dotted lines indicate standard deviation. DMSO, 12 cells; low, 9 cells; medium, 9 cells; high, 10 cells. (D) Recovery curves of mEmerald-NMIIB fluorescence intensity after photobleaching in cells treated with different concentrations of p-nitroblebb. DMSO, 36 cells; low, 36 cells; medium, 30 cells; high, 28 cells. (E) Quantification of the recovery time and the immobile fraction of mEmerald-NMIIB by FRAP. (F) Effect of C3 exoenzyme treatment on the cortical NMIIB and active RhoA in a mitotic cell is shown. The changes of NMIIB and active RhoA intensities at the mitotic cell cortex before and after 10 μg/ml C3 exoenyzme treatments were quantified. The fold change is a ratio of the mean intensity after 10 μg/ml C3 exoenzyme treatment to the mean intensity before treatment. The control group was added with media as C3 exoenzyme was dissolved in water. NMIIB (control), 22 cells; NMIIB (C3 exoenzyme), 37 cells; active RhoA (control), 35 cells; active RhoA (C3 exoenzyme), 36 cells. Data represent means ± SD. Scale bar, 5 μm.

To test the response of active RhoA reporter, mitotic cells were treated with a RhoA inhibitor C3 exoenzyme that prevents activation of RhoA, both active RhoA and NMIIB disappeared from the mitotic cell cortex (Fig. 1 F). This indicates that mRuby2-AHPH reliably reports the active state of RhoA. Taken together, AHPH-mRuby2 and mEmerald-NMIIB report the status of cellular contractility during mitosis and cytokinesis.

### Mechanoaccumulation of NMIIB and active RhoA at the mitotic cell cortex

To study how NMIIB responds to mechanical forces during mitosis, we compressed the mitotic cells with a soft elastic gel and a weight (Fig. 2 A). When compressive forces were exerted on the mitotic cells, the cell has enlarged its diameter, and the cell height has decreased from 22.89 ± 1.88 μm to 15.19 ± 1.19 μm (Figs. 2 B and C). Within 2 minutes after compression, we observed increased intensity of NMIIB at the mitotic cell cortex. In average, there was 1.45 ± 0.29-fold increase of cortical NMIIB intensity in individual cells (Fig. 2 D). Concomitantly, active RhoA increased its intensity at the mitotic cell cortex by 1.54 ± 0.31-fold compared to that of before compression (Figs. 2 E and F). Approximately 76.5% individual cells showed an increase of both active RhoA and NMIIB at the mitotic cell cortex after compression (Fig. S1 A), suggesting that in response to compressive forces, mitotic cells accumulate more active RhoA leading to an accumulation of NMIIB at the cell cortex. Consistently, individual mitotic cells expressing active RhoA reporter using Rhotekin Rho binding domain (Mahlandt et al., 2021) also showed mechanoaccumulation of active RhoA at the cell cortex in response to compressive forces (Fig. S1 B). Furthermore, compressive forces have resulted in more activated NMII as revealed by higher level of phosphorylated myosin regulatory light chain (p-MRLC) in the cell lysate from compressed cells (Fig S1 C).

**Figure 2.**
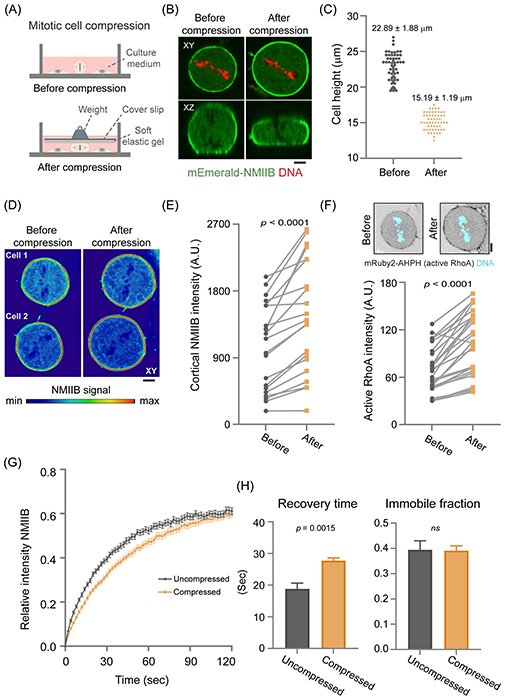
NMIIB and active RhoA show mechanoaccumulation at the cell cortex during mitosis. (A) A schematic representation of the mitotic cell compression experiment. A soft elastic polyacrylamide gel (2 kPa) was used in the study. (B) A representative cell before and after compression is shown. XY, top view; XZ, side view. (C) Quantification of the cell height before and after compression. 56 cells were counted for their cell height before and after compression. (D) Two representative cells with increased accumulation of NMIIB at the cell cortex are shown. (E) Intensity of the cortical NMIIB in mitotic cells before and after compression was quantitated. Black dots indicate the intensity before compression and orange dots indicate the intensity after compression of the same cells. 21 cells were counted. (F) Intensity of active RhoA in mitotic cells before and after compression was quantitated. Black dots indicate the intensity before compression and orange dots indicate the intensity after compression of the same cells. A representative cell with increased cortical active RhoA after compression is shown. 26 cells were counted. (G) Recovery curves of the cortical mEmerald-NMIIB intensity in uncompressed cells and in compressed cells after photobleaching. Uncompressed, 55 cells; compressed, 48 cells. (H) Quantification of the recovery time and the immobile fraction of mEmerald-NMIIB by FRAP. Data represent means ± SD, except for (G) where data represent ± SEM. Scale bar, 5 μm.

To test whether mechanoaccumulation of active RhoA at the cell cortex depends on cell surface tension, we decreased cell surface tension by inhibiting NMII function with p-nitroblebb treatment. We first measured cell surface tension in mitotic cells treated with p-nitroblebb using the micropipette aspiration technique. The mitotic cell surface tension decreased to 56%, 44%, 31% of the control group after treatments with low, medium and high concentrations of p-nitroblebb, respectively (Fig. S1 D; DMSO: 894.90 ± 186.74 pN/μm; [Low]: 500.42 ± 90.64 pN/μm; [Medium]: 393.87 ± 121.08 pN/μm; [High]: 281.58 ± 60.84 pN/μm). Upon p-nitroblebb treatment, we observed that the cortical localization of active RhoA decreased when cell surface tension has been lowered for 3-fold in mitotic cells treated with p-nitroblebb, suggesting that RhoA could respond to the cortical tension generated by NMII during mitosis (Fig. S1 E).

Next, we used FRAP to analyze the cortical NMIIB turnover in mitotic cells under compression. We found that compression in mitotic cells has slowed down the turnover of the cortical NMIIB, with an increase of the recovery half time of 27.75 ± 0.86 s as compared to that of in the uncompressed mitotic cells that was 18.85 ± 1.79 s (Figs. 2 G and H). Thus, compressive forces stabilize NMIIB at the mitotic cell cortex leading to mechanoaccumulation of NMIIB in a RhoA-dependent manner.

### Roles of RhoA/ROCK and CDK1 in mitotic NMIIB mechanoaccumulation under compression

When cells are entering mitosis from G2 phase, cyclin-dependent kinase 1 (CDK1) regulates RhoGEF Ect2 to activate RhoA, which in turn exerts its effects on the actomyosin network through ROCK signaling (Fig. 3 A) (Matthews et al., 2012; Rosa et al., 2015). Since we found that active RhoA responds to compressive forces during mitosis and leads to NMIIB activation and accumulation at the cell cortex, we asked whether CDK1 and ROCK are involved in the mechanoaccumulation of RhoA and NMIIB during mitosis. We first examined the effects of CDK1 and ROCK inhibition in mitotic cells not subjected to compressive forces. When mitotic cells were chemically inhibited by the ROCK inhibitor Y-27632, NMIIB decreased about 2-fold at the cell cortex while active RhoA did not show significant changes, which is consistent with NMIIB functions downstream of ROCK following RhoA activation (Figs. S2 A and B). Consistently, there was about 20% decrease of cell surface tension from 965.34 ± 133.34 pN/μm in the control group to 770.04 ± 160.08 pN/μm in the Y-27632 treated group (Fig. S2 C). Of note, a decrease of cell surface tension by the concentration of Y-27632 used in this study was not sufficient to reduce active RhoA from the cell cortex. When we treated mitotic cells with CDK1 inhibitor RO-3306, NMIIB decreased about 2-fold at the cell cortex and the cell surface tension decreased 15% to 821.67 ± 106.14 pN/μm compared to the control group (Figs. S2 B and C). Interestingly, active RhoA showed an increased accumulation at the cell cortex upon inhibition of CDK1 (Fig. S2 B). Thus, CDK1 appears to regulate NMIIB independent of RhoA activation in mitotic cells, which is different from that of in interphase cells entering mitosis.

**Figure 3.**
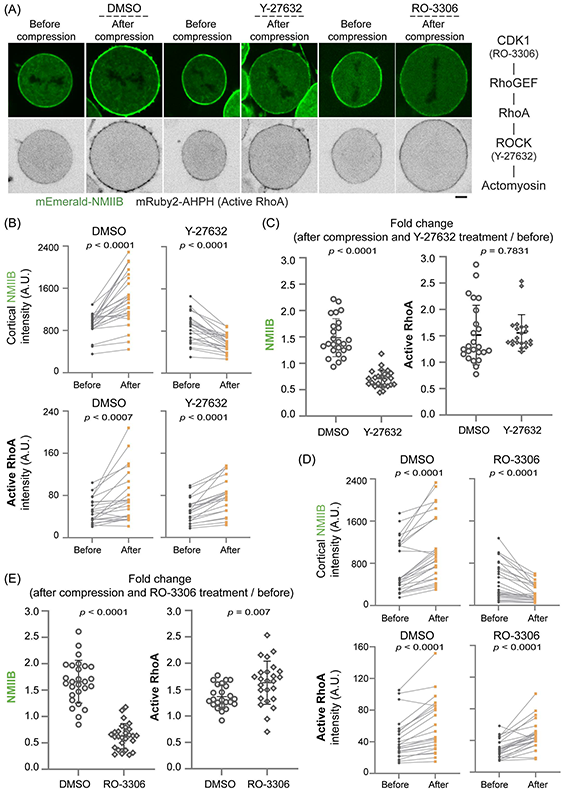
Regulation of NMIIB mechanoresponse by ROCK and CDK1 signaling pathways. (A) The cortical NMIIB and active RhoA at the cell cortex of mitotic cells before and after DMSO or 20 μM Y-27632 treatments or 10 μM RO-3306 are shown. (B) Quantification of the changes of NMIIB and active RhoA intensities at the cell cortex before and after DMSO or 20 μM Y-27632 treatments. (C) The fold change is a ratio of the mean intensity after treatments to the mean intensity before treatment. A ratio higher than 1 means a positive mechanoresponse. NMIIB (DMSO), 26 cells; NMIIB (Y-27632), 47 cells; active RhoA (DMSO), 30 cells; active RhoA (Y-27632), 55 cells. (D) Quantification of the changes of NMIIB and active RhoA intensities at the cell cortex before and after DMSO or 10 μM RO-3306 treatments. (E) The fold change is a ratio of the mean intensity after treatments to the mean intensity before treatment. A ratio higher than 1 means a positive mechanoresponse. NMIIB (DMSO), 27 cells; NMIIB (RO-3306), 27 cells; active RhoA (DMSO), 24 cells; active RhoA (RO-3306), 25 cells. Data represent means ± SD. Scale bar, 5 μm.

Next, we examined the mechanoresponse of NMIIB and active RhoA in mitotic cells treated with ROCK or CDK1 inhibitors (Fig. S2 D). When mitotic cells were compressed for about 8 minutes in the presence of Y-27632, there was a decrease of cortical NMIIB as compared to the control group that showed an increase of cortical NMIIB after compression (Figs. 3 A, B and C). Active RhoA accumulated at the mitotic cell cortex after compression in the presence or absence of Y-27632 (Figs. 3 A and B and C). Similarly, when mitotic cells were compressed in the presence of RO-3306, NMIIB decreased its intensity at the cell cortex while active RhoA showed mechanoaccumulation at the cell cortex (Figs. A D and E). Collectively, our data show that ROCK and CDK1 are required for NMIIB to accumulate at the cell cortex in response to compressive forces during mitosis, however, CDK1 appears to exert its effect on NMIIB independent of active RhoA.

### Phosphoregulation of NMIIB tail region is not involved in NMIIB mechanoaccumulation

RhoA / ROCK signaling modulates NMII through phosphorylation of MRLC that associates with NMII’s head domain. In addition, NMII is also regulated by the phosphorylation of its tail region, which is involved in regulating myosin II filament assembly (Heissler and Sellers, 2016). Phosphorylation of NMIIB S1935 lowers the ability of NMIIB to assemble into myofilaments (Fig. 4 A) (Juanes-Garcia et al., 2015; Schiffhauer et al., 2019). To study the role of NMIIB tail phosphorylation, we first generated three cell lines stably expressing wild type NMIIB, NMIIB containing a non-phosphorylatable alanine residue at S1935 (S1935A), and NMIIB containing a phosphomimetic aspartate residue at S1935 (S1935D), respectively. The endogenous NMIIB in these cell lines was depleted using CRISPR interference (CRISPRi) that switches off gene expression from its promoter and does not interfere with the transgene expression (Fig. S3 A and B). Cells expressing NMIIB S1935D showed a decrease in the cytoskeletal fraction compared to cells expressing wild type NMIIB or NMIIB S1935A, indicating less cytoskeletal-bound NMIIB myofilaments were present (Fig. 4 B and C).

**Figure 4.**
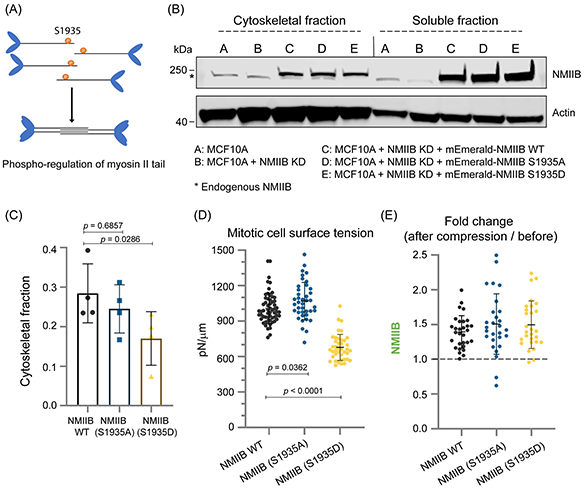
Perturbation of the NMIIB tail domain regulation affects maintenance of the mitotic cell surface tension but less on the NMIIB mechanoresponse. (A) A schematic representation of the phosphoregulation of the NMIIB tail regrion in controlling the myosin II filament assembly. (B) A protein blot shows the cytoskeletal fractions and soluble fractions of NMIIB and actin in MCF10A cells expressing different phosphomutants in the absence of endogenous NMIIB. KD, knockdown. (C) Quantification of the cytoskeletal fractions of NMIIB wild type or phosphomutants from the protein blot. (D) Measurement of the mitotic cell surface tension by MPA in cells expressing NMIIB (WT), NMIIB (S1935A), or NMIIB (S1935D). NMIIB (WT), 59 cells; NMIIB (S1935A), 40 cells; NMIIB(S1935D), 40 cells. (E) Quantification of the changes of NMIIB (WT), NMIIB (S1935A), and NMIIB (S1935D) at the cell cortex before and after compression. The fold change is a ratio of the mean intensity after compression to the mean intensity before compression. The dotted indicates ratio of 1 in which a ratio higher than 1 means a positive mechanoresponse. NMIIB (WT), 30 cells; NMIIB (S1935A), 28 cells; NMIIB (S1935D), 28 cells. Data represent means ± SD. Scale bar, 5 μm.

When we examined the NMIIB tail phosphomutants, we found that mitotic cells expressing NMIIB S1935D generated lower cell surface tension (1007.94 ± 130.21 pN/μm in wild type NMIIB and 677.67 ± 111.03 pN/μm in NMIIB S1935D) but were able to demonstrate mechanoaccumulation of NMIIB S1935D after compression (Figs. 4 D and E; S3 C). Mitotic cells expressing NMIIB S1935A exhibited a similar level of cell surface tension (1069.76 ± 158.13 pN/μm in NMIIB S1935A) and mechanoaccumulation as in cells expressing wild type NMIIB (Figs. 4 D and E; S3 C). Thus, phosphorylation of NMIIB S1935 at the tail domain affected NMIIB’s function in cell surface tension generation but less on the mechanoresponse of NMIIB, in line with the previous study showing that the NMIIB tail domain fine-tuned the NMIIB mechanosensitive dynamics and the phosphomutants could accumulate with varying degree at the cell cortex in epithelial cells in response to mechanical stress (Schiffhauer et al., 2019).

### NMIIB translocation activity is required for its mechanoaccumulation under compression

Our results showed that regulation of NMII’s head domain is important for NMIIB to accumulate at the cell cortex in response to compressive forces during mitosis. NMII’s head domain has two properties namely actin-binding functions and motor activity to translocate actin filaments (Sweeney and Houdusse, 2010). Previous studies showed that NMII’s mutations that selectively uncouple these two properties have revealed distinct contributions of each property to NMII’s cellular functions (Choi et al., 2008; Ma et al., 2012; Smutny et al., 2010). To see the roles of these properties in regulating NMIIB mechanoaccumulation, we expressed two previously characterised NMIIB mutants NMIIB R709C or NMIIB R241A that are defective in motor activity but retaining their ability to bind actin filaments (Kim et al., 2005; Osorio et al., 2019), and one NMIIB mutant K647A that has a weakening actin-binding function (von der Ecken et al., 2016) in the reporter cells (Fig. 5 A). We further depleted the endogenous NMIIB in these cell lines using CRISPRi.

**Figure 5.**
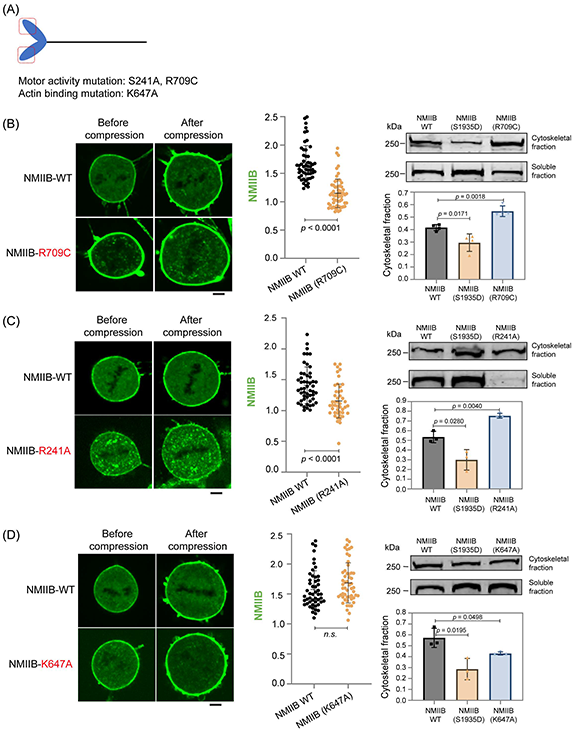
NMIIB that is motor-dead but retaining actin-binding function is defective in mechanoaccumulation. (A) A diagram showing the motor activity mutations and actin-binding mutations of NMIIB used in the study. (B) Left panel, micrographs of a cell expressing wild type NMIIB (NMIIB-WT) and a cell expressing NMIIB-R709C before and after compression; middle panel, the fold change is a ratio of the mean intensity after compression to the mean intensity before compression. A ratio higher than 1 means a positive mechanoresponse.NMIIB WT, 53 cells; NMIIB (R709C), 60 cells; right panel, a protein blot and quantification of the cytoskeletal fractions and soluble fractions of NMIIB wild type and NMIIB (R709C) mutant proteins. (C) Left panel, micrographs of a cell expressing wild type NMIIB (NMIIB-WT) and a cell expressing NMIIB-R241A before and after compression; middle panel, the fold change is a ratio of the mean intensity after compression to the mean intensity before compression. A ratio higher than 1 means a positive mechanoresponse. NMIIB WT, 51 cells; NMIIB (R241A), 46 cells; right panel, a protein blot and quantification of the cytoskeletal fractions and soluble fractions of NMIIB wild type and NMIIB (R241A) mutant proteins. (D) Left panel, micrographs of a cell expressing wild type NMIIB (NMIIB-WT) and a cell expressing NMIIB-K647A before and after compression; middle panel, the fold change is a ratio of the mean intensity after compression to the mean intensity before compression. A ratio higher than 1 means a positive mechanoresponse. NMIIB WT, 50 cells; NMIIB (K647A), 54 cells; right panel, a protein blot and quantification of the cytoskeletal fractions and soluble fractions of NMIIB wild type and NMIIB (K647A) mutant proteins. Data represent means ± SD. Scale bar, 5 μm.

Cytoskeletal fractionation analyses showed that cells expressing NMIIB R709C or R241A have an increased NMIIB cytoskeletal fraction than cells expressing wild type NMIIB or NMIIB S1935D, indicating a strong cytoskeletal binding of these two NMIIB mutant proteins despite a defective motor activity (Figs. 5 B and C, right panels). Interestingly, when we compressed the mitotic cells expressing NMIIB R709C or R241A, there was no significant increase of cortical NMIIB intensity compared to that of in control cells expressing wild type NMIIB (Figs. 5 B and C, left and middle panels). In contrast, compression of mitotic cells expressing NMIIB K647A, which has more NMIIB mutant proteins in the soluble fraction, showed a similar mechanoaccumulation of NMIIB at the cell cortex as in control cells. Thus, our data show that the translocation activity of NMIIB, but not just the actin-binding function of the NMIIB’s head domain, is required for NMIIB to accumulate at the cell cortex in response to compressive forces during mitosis.

### NMIIB translocation activity has a dominant role in regulating its mechanoaccumulation

Next, we asked whether the NMIIB tail mutation S1935D could reduce the tight cytoskeletal association of the NMIIB translocation mutants and hence reverting NMIIB mechanoresponse. To this end, we constructed three NMIIB mutants containing double mutations R709C S1935D, R241A S1935D and K647A S1935D, respectively. We first determined the proportion of the cytoskeletal fraction in protein lysates extracted from these NMIIB double mutants. The results showed that cells expressing NMIIB R709C S1935D and NMIIB R241A S1935D have more insoluble cytoskeletal-bound NMIIB mutant proteins compared to control cells (Fig. 6 A). Compression of these two NMIIB mutant cells showed no significant increase of the cortical NMIIB at the cell cortex compared to cells expressing wild type NMIIB or NMIIB S1935D (Figs. 6 B and C). Cells expressing NMIIB K647A S1945D, have more mutant proteins in the soluble fraction compared to control cells and retain the ability to accumulate the cortical NMIIB double mutant protein at the cell cortex in response to compression (Figs. 6 A B and C). Our data show that NMIIB S1935D mutation does not revert mechanoaccumulation defects of NMIIB translocation mutants, and the NMIIB translocation activity at its head domain plays a dominating role in regulating NMIIB mechanoaccumulation at the cell cortex during mitosis.

**Figure 6.**
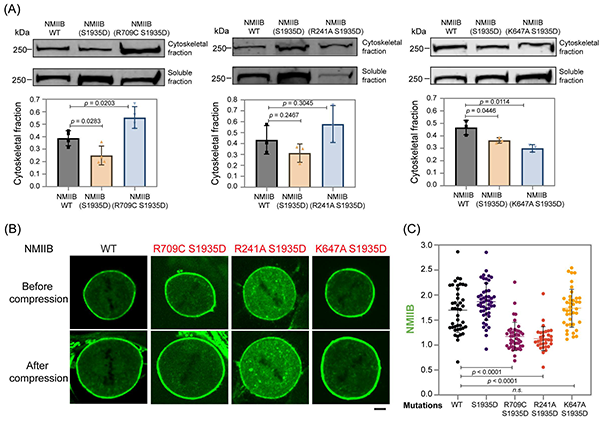
The translocation activity plays a dominant role in regulating NMIIB mechanoaccumulation. (A) Protein blots and quantification of the cytoskeletal fractions and soluble fractions of NMIIB wild type and NMIIB double mutant proteins. (B) Micrographs of cells expressing wild type NMIIB or NMIIB double mutants before and after compression. (C) The fold change is a ratio of the mean intensity after compression to the mean intensity before compression. A ratio higher than 1 means a positive mechanoresponse. NMIIB WT, 43 cells; NMIIB (S1935D), 48 cells; NMIIB (R709C S1935D), 43 cells; NMIIB (R241A S1935D), 32 cells; NMIIB (K647A S1935D), 40 cells. Data represent means ± SD. Scale bar, 5 μm.

## Discussion

NMIIs accumulate at the subcellular region with applied mechanical stresses (Luo et al., 2013). Mechanoresponse of NMIIs facilitates cellular mechanosensing during cell migration and cell division and confers mechanical adaptation to cancer cells living in physically challenging microenvironment (Nguyen et al., 2022). Within a tissue environment, cells constantly experience mechanical stresses such as compressive stress from surrounding cells and the extracellular matrix. This is particularly significant in an overgrowth tumor environment where solid stress or compressive stress accumulates inside the tumors (Nia et al., 2020). Normal or cancer cells that are not able to mechanically adapt to the constantly changing microenvironment by engaging mechanoresponse accumulate cell division defects that lead to aneuploidy (Cattin et al., 2015; Matthews et al., 2020).

During the cross-bridge cycle of force generation, NMII binds and slides actin filaments (Sweeney and Houdusse, 2010). Interestingly, these two closely related biophysical properties can be genetically uncoupled in different cellular functions of NMIIs. Previous studies showed that maturation of nascent adhesions in migrating cells and division of certain cell types require NMIIs to bind and crosslink actin networks but does not involve its motor activity (Choi et al., 2008; Ma et al., 2012). In contrast, NMIIs bind and slide actin filaments during cell-cell adhesions and embryonic cell division (Kasza et al., 2019; Osorio et al., 2019; Smutny et al., 2010). We tested this differential requirement of NMII in its mechanoresponse during mitosis. Using two NMIIB mutations (R709C and R241A in NMIIB) that are motor dead but retaining actin-binding function (Ma et al., 2012; Osorio et al., 2019), we revealed that mechanoaccumulation of NMIIB at the mitotic cell cortex requires the translocation activity of NMIIB. Interestingly, a mutation in loop 2 of the actin-binding surface of NMIIB (K647A in NMIIB), which regulates the cleft closure of U50 and L50 subdomains of NMIIs for strong actin-binding function (Onishi et al., 2006; von der Ecken et al., 2016), does not dampen NMIIB mechanoresponse. Thus, actin-binding function of NMIIs alone is not sufficient to regulate mechanoaccumulation of NMIIB at the mitotic cell cortex, and counterintuitively, the motor activity of NMIIB in driving actin filament translocation is essential in NMIIB mechanoresponse.

Activation of RhoA/ROCK signaling pathway leads to MRLC phosphorylation and is associated with an increase of the actin-activated ATPase kinetics of myosin head domain and hence its motor activity (Amano et al., 1996; Heissler and Sellers, 2016; Nayak et al., 2020). We found that mechanoaccumulation of NMIIB is a result of active RhoA mechanoaccumulation at the mitotic cell cortex. Through the effector ROCK, mechanoaccumulation of active RhoA drives higher activation of myosin motor domain and stabilization of NMIIB at the cell cortex in response to compressive forces in mitotic cells. Interestingly, NMIIB mechanoaccumulation at the cell cortex can be independent of active RhoA accumulation as CDK1 inhibition leads to decreased cortical NMIIB but higher cortical active RhoA in compressed mitotic cells. Similar observation has been identified in starfish oocytes undergoing meiotic division in which RhoA is activated when CDK1 is inhibited during surface contraction waves (Bischof et al., 2017). Thus, CDK1 regulates NMIIB mechanoaccumulation during mitosis independent of active RhoA. This contrasts with CDK1’s function in activating Ect2/RhoA/ROCK signaling and NMIIs when cells enter mitosis from interphase (Matthews et al., 2012; Rosa et al., 2015).

Mechanical activation of RhoA happens at the cell-cell adherens junctions and the cell-matrix focal adhesion to reinforce actomyosin contractility in response to mechanical stress (Acharya et al., 2018; Guilluy et al., 2011). Our study, to our knowledge, is the first to show accumulation of active RhoA at the cell cortex in response to compressive forces during mitosis. Mechanoaccumulation of active RhoA at the cell cortex during mitosis was not previously identified although inhibition of its downstream effector ROCK led to a decreased mechanoresponse of NMII (Schiffhauer et al., 2016). In our study, we employed widely used live-cell active RhoA reporters (Koh et al., 2021; Mahlandt et al., 2021) and focused only on mitotic cells using a microscopy with confocality and high temporal resolution. This has facilitated a sensitive detection of RhoA mechanoresponse at the cell cortex when mitosis is mechanically challenged.

Our finding implies that NMII mechanoaccumulation involves lever arm movement generated by the motor activity. Lever arm movement is coupled to nucleotide state changes during Pi release and ADP release and generates NMIIs of different nucleotide states and actin-binding affinity (Debold, 2021; Sweeney and Houdusse, 2010; Trivedi et al., 2015; Wulf et al., 2016). Consistently, NMIIs engineered to have longer neck structure and hence larger lever arm movement have stronger ability to mechanoaccumulate in cells (Luo et al., 2013). We propose a model that NMIIB species of different actin-binding affinity are generated after lever arm movement and nucleotide state changes. These NMIIB species bind actin filaments at varying affinity and in ensemble display accumulation in response to compressive forces during mitosis. This notion is supported by the existence of “Goldilocks zone” for the actin-binding affinity for an optimal mechanoaccumulation as described for actin-binding proteins such as α-actinin, filamin and NMIIs (Schiffhauer et al., 2016).

## Materials and methods

### Cell lines and culture

MCF10A cell lines were cultured in DMEM F-12 (Shanghai BasalMedia) with Glutamax (Gibco), 5% horse serum (Biological Industries), 20 ng/ml EGF (Gibco), 0.5 mg/ml Hydrocortisone (MedChemExpress), 100 ng/ml Cholera toxin (Sigma), 10 μg/ml Insulin (Biological Industries), 1% Pen/Strep (Biological Industries) at 37°C with 5% CO2. The MCF10A cell line was a gift from Dr. Kuan Yoow Chan from Zhejiang University. The cells were authenticated by Sangon Biotech and were routinely checked for mycoplasma contamination in the lab using Myco-Blue Mycoplasma Detector (Vazyme). MCF10A cells expressing mRuby2-AHPH and mEmerald-NMIIB, or dTomato-2xrGBD were prepared by lentivirus transduction and were enriched using the fluorescence-activated cell sorting (FACS). MCF10A cells expressing mEmerald-NMIIB WT, mEmerald-NMIIB S1935A, and mEmerald-NMIIB S1935D, mEmerald-NMIIB R709C, mEmerald-NMIIB R241A, mEmerald-NMIIB K647A, and all NMIIB double mutants, respectively, were prepared by lentivirus transduction and selected with 2 μg/ml puromycin (Sangon Biotech) and were enriched using the FACS. To knockdown the endogenous NMIIB, the lentivirus expressing pCRISPRi0003 containing the guiding RNA (gRNA) targeting the NMIIB’s 5’ UTR region was transduced into the cells expressing NMIIB mutants. Cells were selected with 6 μg/ml blasticidin (Solarbio) for 6 days and were continued to grow without the antibiotic for another 1 to 3 days before the experiments.

### Drug treatments

The following drugs were used in the study: NMII inhibitor: p-nitro-blebbistatin (Cayman Chemical, 24171); Rho inhibitor: C3 exoenzyme (Cytoskeleton, CT04); ROCK inhibitor: Y-27632 (MedChemExpress, HY-10583); CDK1 inhibitor: RO-3306 (Selleckchem, S7747). DMSO was used to dissolve p-nitro-blebbistatin, Y-27632 and RO-3306. C3 exoenzyme was dissolved in water. Details of concentrations used in the study are described in figure legends.

### Lentiviral expression constructs

The following lentiviral transfer plasmids were used in the study: pTGL0427 containing mRuby2-AHPH in which AHPH was subcloned from an Addgene plasmid #68026 (a gift from Michael Glozter) and fused with mRuby2; pTGL0193 containing mEmerald-NMIIB, which was subcloned from an Addgene plasmid #54192 (a gift from Michael Davidson); pTGL0594 containing dTomato-2xrGBD subcloned from an Addgene plasmid #129625 (a gift from Dorus Gadella); pTGL0373 containing mEmerald-NMIIB S1935A; pTGL0374 containing mEmerald-NMIIB S1935D; pTGL0584 containing mEmerald-NMIIB R709C; pTGL0585 containing mEmerald-NMIIB R709C S1935D; pTGL0582 containing mEmerald-NMIIB R241A; pTGL0583 containing mEmerald-NMIIB R241A S1935D; pTGL0589 containing mEmerald-NMIIB K647A; pTGL0590 containing mEmerald-NMIIB K647A S1935D; pTGL0386 containing KRAB and dCas9 for CRISPRi (a gift from Jorge Ferrer, Addgene plasmid #118154); pCRISPRi0001 containing gRNA 5’ GTGCTAAAGGAGCCCGGCGG 3’ cloned into pTGL0386; pCRISPRi0002 containing gRNA 5’ GCTGGATCTGTGGTCGCGGC 3’ cloned into pTGL0386; pCRISPRi0003 containing gRNA 5’ GGACTGAGGCGCTGGATCTG 3’ cloned into pTGL0386. The 3^rd^ generation lentiviral packaging system was used to prepare lentiviral particles. In brief, packaging plasmids pRSV-Rev, pMDLg/pRRE, pMD2.G (gifts from Didier Trono) were chemically transfected with the transfer plasmid into the 293Ta packaging cell line (Genecopoeia, LT008) using GeneTwin transfection reagent (Biomed, TG101). After 3 days, the culture medium was collected, filtered through a 0.45 μm filter, and then concentrated using the lentivirus concentration solution (Genomeditech, GM-040801) before adding to MCF10A cells.

### Mitotic cell preparation

To increase the mitotic cell number, cells cultured on the imaging chambered coverglass were treated with 7.5 μM RO-3306 (Selleckchem) for 16 to 18 hours at 37 °C and were washed with the pre-warmed culture medium for 3 times to release cells into mitosis and added with 1 ml of fresh medium. Experiments were performed 40 minutes after the drug wash-off.

### Elastic polyacrylamide gel preparation

The polyacrylamide gel used for cell compression was prepared as described in Matthews *et al*. and Le Berre *et al*. with modifications (Le Berre et al., 2014; Matthews et al., 2020). Briefly, 18 mm glass coverslips (Sangon, F518211) were treated with 10 μl Binding-silane (Sangon, C500226) for 10 minutes and then were rinsed with 100% ethanol and air-dried. To prepare soft elastic gels (2 kPa), 1 ml polyacrylamide gel solution was prepared by mixing 125 μl 40% w/v acrylamide (Sangon), 35 μl 2% bis-arcylamide (Sangon), 10 μl APS (10% in water, Sangon), and 830 μl water. After adding and mixing 1 μl TEMED (Sangon) into the gel solution, about 350 μl of the final gel solution was immediately transferred onto a flat glass slide and covered by the coverslip pre-treated with the Binding-silane. After polymerization for 20 minutes, the gel and the attached coverslip were gently removed from the glass slide using a surgical blade and were soaked in PBS for at least 2 hours and followed by incubation in cell culture media for overnight. In experiments that involved drug treatment, the gels were incubated in media containing the drugs for overnight.

### Micropipette aspiration (MPA)

The micropipette was prepared by first pulling a borosilicate glass capillary (Sutter Instrument, B100-58-10) using the P-97 Micropipette puller (Sutter Instrument). Then, the thin end of a borosilicate glass capillary was cut by a Microforge (Narishige) to a diameter of about 5.5 μm. The micropipette was bent to a desired degree by heating it over an alcohol lamp and was filled with phosphate-buffered saline (PBS) using the MicroFil (World Precision Instruments, MF34G-5). The micropipette was installed to a micropipette holder, which was connected to a syringe pump (Harvard apparatus) and a 25 ml serological plastic pipettes via a three-way valve. The micropipette was positioned under an inverted microscope (Olympus, IX73) equipped with Olympus 60x objective lens (N.A. 1.35, U Plan super apochromat), a micromanipulator system (Eppendorf, TransferMan 4r), and a CMOS high speed camera (Vision Research, Phantom 410L, 333.33 nm / pixel).

To perform micropipette aspiration, cells were cultured on a one-well chambered coverglass (Cellvis, C1-1.5H-N) for 24 hours and then were treated with 7.5 μM RO-3306 for 18 hours. Cells were washed 3 times with fresh media and were added with fresh media containing 20 mM HEPES. Cells were incubated for another 40 minutes to reach mitosis. When a mitotic cell was aspirated by the micropipette, the microscopy image was captured and analyzed using Fiji to obtain the pipette radius (R_p_) and the cell radius (R_c_). The cell length of about half a diameter of the pipette opening was aspirated in. The water pressure changes (ΔP) as displayed on the 25 ml serological plastic pipette was also recorded. Cell surface tension (T) was then calculated based on the Laplace equation, T = ΔP / 2 (1/R_p_ – 1/R_c_).

### Spinning-disk confocal microscopy for live-cell imaging

Spinning-disk confocal microscopy was equipped with a Nikon Eclipse Ti2-E inverted microscope, Nikon 60x oil-immersion objective lens (N.A. 1.40, Plan Apochromat Lambda), a spinning-disk system (Yokogawa Electric Corporation, CSU-W1), a Photometrics Prime 95B sCMOS camera, a Piezo Z stage (Physik Instrumente), and a live-cell stage top chamber with humidified CO_2_ (Okolab). Images were acquired using the Metamorph with a z-step size of 0.5 μm and a x-y plane resolution of 183.33 nm / pixel. The fluorophores were excited by laser lines at wavelengths of 488, 561, or 640 nm. For experiments involving cell compression, a z-stack of 30 μm was acquired before cells were compressed and a stack of 22 μm was acquired after cells were compressed. This ensured that the entire cell volume was covered during imaging.

### Mitotic cell compression and live-cell imaging

To compress cells, 8 x 10^4^ cells were first seeded on a 20 mm two-well chambered cover glass (Cellvis, C2-1.5H-N). Forty-eight hours later, cells were treated with 7.5 μM RO-3306 to synchronize cells at the G2/M boundary for 16 to 18 hours. Prior to live-cell imaging with compression, synchronized cells were washed 3 times with the pre-warmed medium and added with 1 ml of fresh medium and incubated in the stage-top chamber attached to the spinning-disk microscopy for 40 minutes. When compressing the mitotic cells, the coverslip coated with the elastic gel was put over the cell layer and followed by a 5 g weight. Elastic gels and weights were pre-heated to 37 °C in the chamber. In experiments where cells were treated with DMSO, 20 μM Y-27632 or 10 μM RO-3306, 1 ml medium containing 40 μM Y-27632 or 20 μM RO-3306 or containing equivalent volume of DMSO was added to cells grown in 1 ml medium on the chambered coverglass to achieve a final concentration of 20 μM Y-27632 or 10 μM RO-3306. The elastic gel that was pre-incubated with 20 μM Y-27632 or 10 μM RO-3306 for overnight was then put on top of the cell layer and followed by a 5 g weight. Cells at the same position were imaged before and after compression. The cell compression efficiency was validated by their decrease of cell heights. For microscopy imaging not involving cell compression, the μ-Slide 8-well chambered coverglass (ibidi, 80826) was used for cell culture and imaging. Mitotic chromosomes were labeled with 0.2 μM SiR-DNA in live cells during imaging (Cytoskeleton, CY-SC007).

### Image analysis and processing

Microscopy images were analyzed using Fiji. To quantitate fluorescence intensity of mEmerald-NMIIB, mRuby2-AHPH and dTomato-2xrGBD in mitotic cells (Figs. 1 C, 1 F, S1 E, S2 A) and in mitotic cells for compression experiments (Figs. 2 E, 2 F, 3 B, 3 C, 3 D, 3 E, 4 E, 5 B, 5C, 5D, 6C), image stacks containing 5 slices around the mid-plane were projected along the z-axis using the sum-intensity projection (Fiji/image/stacks/Z project). Background subtraction (Fiji/process/subtract background) was performed on projected images using the rolling ball radius of 20 pixels. Then, a segmented line with a line width of 4 (for mitotic cells) or 5 (for mitotic cells in compression experiments) was used to select the cortical fluorescent signals over the entire cell circumference. The mean fluorescence intensity of the selected line region from the same cells before and after treatment and/or compression was measured. The fold change was then calculated by dividing the intensity after treatment and/or compression to the intensity before treatment and/or compression. For time series in Fig. 1 B, the mean intensity of the cortical fluorescence was measured every 45 s. To quantitate the cell height in cell compression experiments, the number of slices from top to bottom of a cell were multiplied by 0.5 μm.

### FRAP

FRAP was performed using the above-mentioned spinning-disk confocal microscopy equipped with Nikon 100x oil-immersion objective lens (N.A. 1.45, Plan Apochromat Lambda) and iLAS2 FRAP system (Gataca). Images were acquired using the Metamorph software at 110 nm / pixel. The mEmerald-NMIIB fluorescence signal was photobleached by using a short pulse of high power 488 nm laser lines and imaged by using a low power 488 nm laser lines. A small bleaching rectangular region-of-interest (ROI) of about 2 x 10-μm was drawn on the cortical mEmerald-NMIIB in mitotic cells. Three frames of pre-bleached images were acquired with 1 second interval before photobleaching, followed by time-lapse imaging of the same cells with 2 seconds interval for 120 seconds. The bleaching ROIs were recorded to locate the regions of bleaching during analysis.

Fiji was used to quantitate the fluorescence intensity of FRAP images. Background subtraction (Fiji/process/subtract background) was performed on the images with the rolling ball radius of 20 pixels. A segmented line ROI (line width 5) was used to select the FRAP regions for quantification. The mean intensity of ROIs for pre-bleached images and after bleached images (I_bleached_) was calculated for each time point. A mean pre-bleached intensity (I_prebleach_) was derived from the average of three pre-bleached images. The ratio of I_bleached_ / I_prebleach_ at each time point was calculated and was subsequently deducted by the ratio of the first image after photobleaching to obtain ratio_t_. The ratio_t_ of each time point was plotted as the recovery curve and fitted using a non-linear fitting function (one-phase association function constrained by Y0 = 0 and plateau < 1) in the Prism 9 software (GraphPad). The recovery time was derived from the half-time parameter calculated from the fitting. The mobile fraction was derived from the span parameter calculated from the fitting. The immobile fraction was calculated as a difference between 1 and the mobile fraction.

### Cytoskeletal fractionation

Cells were washed once with ice-cold PBS. To a single well of a 6-well cell culture plate, 200 μl of lysis buffer (50 mM PIPES, pH 6.8, 46 mM NaCl, 2.5 mM EGTA, 1 mM MgCl_2_, 1mM ATP, 0.5% Triton-X 100, protease inhibitor cocktail) was added. Lysates were harvested and centrifuged at 13000 *g* for 20 minutes at 4°C. The supernatant (soluble fraction) was separated from the pellet (cytoskeletal fraction). The pellet was then resuspended in a same volume of lysis buffer as the supernatant. Both fractions were added with 5 x Laemli sample buffer and boiled for 5 minutes. The pellet suspension was sonicated in a water bath sonicator for 1 minute with 20% power to improve solubility.

### Protein immunoblotting

Protein samples were separated in a pre-cast SDS-PAGE gel (GenScript, M00657) and blotted to a PVDF membrane (EMD Millipore, immobilon-P, IPVH00010). Blots were blocked with 5% skim milk for 1 hour at room temperature and were incubated at 4°C overnight with the NMIIB antibody (Proteintech, 19673-1-AP; used in Figs. 4 B and S3 A) or the β-actin antibody (HuaBio, ET1701-80; used in Figs. 4 B and S1 C) or the GFP antibody (HuaBio, ET1604-26; used in Figs. 5 B C D and 6 A) or GAPDH antibody (Cell Signaling Technology, #5174; used in Fig. S3 A) or p-MLC2 antibody (Cell Signaling Technology, #3675; used in Fig. S1 C). The secondary antibodies used were Alexa Fluor Plus 800-conjugated goat anti-rabbit IgG secondary antibody (ThermoFisher, A32735) and Alexa Fluor 680-conjugated goat anti-mouse IgG secondary antibody (ThermoFisher, A32729). Blots were imaged using the LI-COR Odyssey CLx imaging system. The signal intensity of protein bands was calculated using the LI-COR Image Studio software and Fiji. The cytoskeletal fraction was calculated by dividing the cytoskeletal fraction intensity by the sum intensity of cytoskeletal and soluble fractions. Protein band intensity measurements in Fig. 4 were obtained from four proteins blots derived from four independent experiments. Protein band intensity measurements in Fig. 5 and 6 were obtained from three proteins blots derived from three independent experiments. The protein lysate in Fig. S1 A was prepared using Minute™ Total Protein Extraction Kit for Animal Cultured Cells and Tissues (Invent Biotech, SD-001). Protein lysates used in protein blots in Figs. 4 and 5 and 6 and S3 were prepared using the cytoskeletal fractionation lysis buffer.

### Quantitative real-time PCR (qPCR) for checking endogenous NMIIB

To validate the knockdown of endogenous NMIIB in cells used in Figs. 4 and 5 and 6, total RNAs were prepared from cells using the FastPure Cell/Tissue Total RNA Isolation Kit (Vazyme, RC101-01) and were reverse transcribed into cDNA using the HiScript II Q RT SuperMix for qPCR (Vazyme, R223-01). The expression level of cDNAs was determined using the real-time PCR with the ChamQ Universal SYBR qPCR Master Mix (Vazyme, Q711-02). The reaction mix was assembled on the hard-shell PCR plate (Bio-Rad, HSP9655), sealed with the Microseal ‘B’ seal (Bio-Rad, MSB1001), and was performed in the CFX96 Touch Real-Time PCR Detection System (Bio-RAD, C1000). Primers used in the qPCR: GAPDH, 5’ CAGGAGGCATTGCTGATGAT 3’ and 5’ GAAGGCTGGGGCTCATTT 3’; NMIIB, 5’ CCTCATGCTGACCTTGCAAA 3’ and 5’ GGACACAAAACCAATATTCCCATT 3’. The primer pair for NMIIB targets the 3’ UTR of NMIIB gene, allowing checking of endogenous NMIIB expression. The C_T_ value for GAPDH was used for normalization to obtain the relative expression level.

### Statistical analysis

Statistical analysis was performed using Prism 9 (GraphPad). Datasets were analyzed by Student t-test. If datasets were not normally distributed, they were analyzed using the Mann-Whitney test. All graphs were plotted using Prism 9 (GraphPad).

## Author contributions

C.W. and J.D. designed, performed experiments, and analyzed data. S.D. prepared elastic gels used in the study. Q.W and X.G provided the devices and technical expertise for micropipette aspiration. T.G.C conceived the study, designed experiments, supervised the study, and prepared the manuscript with inputs from all authors.

## Acknowledgement

This study was funded by Zhejiang University International Campus start-up grant (130000-541902/016) and National Natural Science Foundation of China grant (32270770) given to T.G.C. We thank Dr. Di Chen for sharing the CRISPRi vector with us. We thank Dr. Mikael Bjorklund for sharing the 293Ta packaging cell line with us. We thank Dr. Wanzhong Ge and Dr. Junqi Huang for their comments on the manuscript. Special thanks to Dr. Fang Kong from the Nanyang Technological University, Singapore for designing the imaging chamber adaptor used in the microscopy imaging for cell compression. Q.W and X.G would like to acknowledge the support from the National Natural Science Foundation of China (12072198).

---

**Figure S1.**
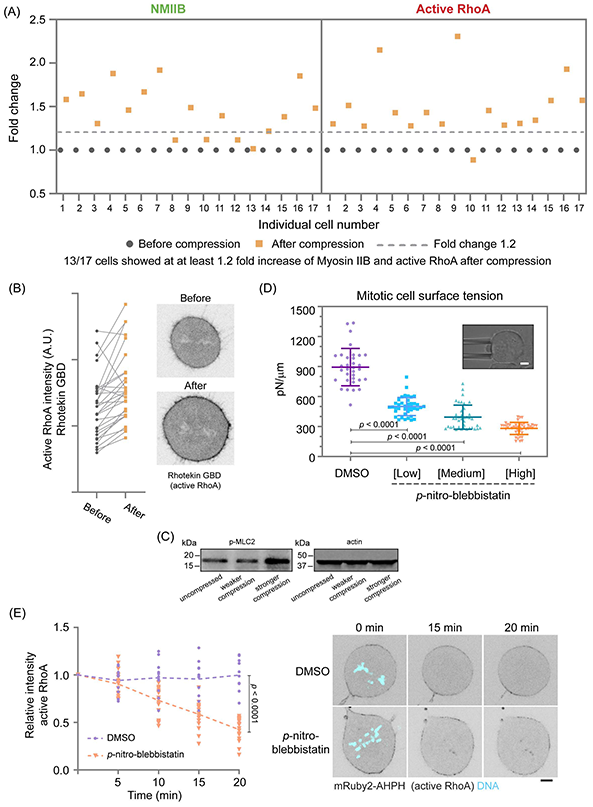
(A) Increased accumulation of NMIIB and active RhoA at the mitotic cell cortex in the same individual cells. Fold changes of NMIIB and active RhoA at the mitotic cell cortex after compression in individual cells were plotted. (B) Intensity of active RhoA in mitotic cells expressing dTomato-2xrGBD (Rhotekin GBD) before and after compression was quantitated. Black dots indicate the intensity before compression and orange dots indicate the intensity after compression of the same cells. A representative cell with increased cortical active RhoA after compression is shown. 29 cells were counted. (C) Protein blots showing levels of phosphorylated myosin light chain and actin in uncompressed and compressed cells. (D) Measurement of the mitotic cell surface tension by MPA in the presence of different concentrations of p-nitroblebb. The inset shows an example of an aspirated cell. DMSO, 33 cells; low, 40 cells; medium, 39 cells; high, 43 cells. (E) Relative changes of active RhoA intensity at the cell cortex after DMSO or 110 μM p-nitroblebb treatments as a function of time. The fluorescence intensity at each time point was normalized to that of in cells before treatments. DMSO, 11 cells; p-nitroblebb, 17 cells. Representative cells showing the decrease of active RhoA at the mitotic cell cortex before and after p-nitroblebb treatment are shown. Data represent means ± SD. Scale bar, 5 μm.

**Figure S2.**
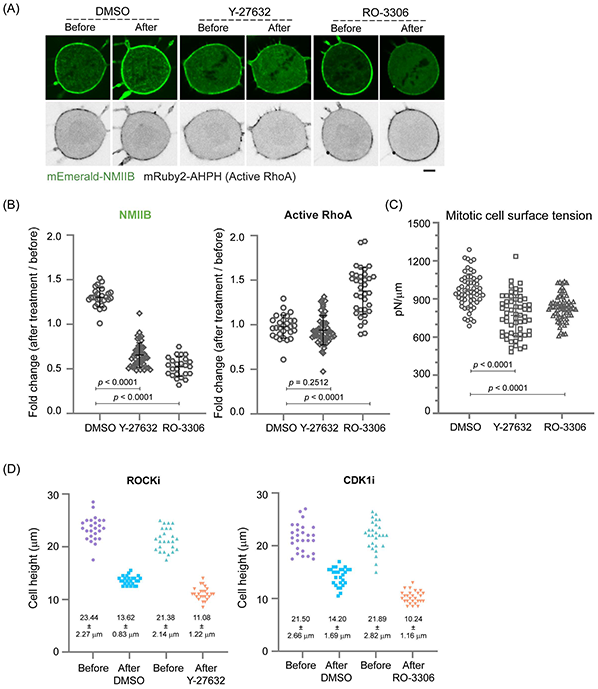
(A) The cortical NMIIB and active RhoA at the cell cortex of mitotic cells before and after DMSO, 20 μM Y-27632 or 10 μM RO-3306 treatments are shown. (B) Quantification of the changes of NMIIB and active RhoA intensities at the cell cortex before and after DMSO or 20 μM Y-27632 treatments. The fold change is a ratio of the mean intensity after treatments to the mean intensity before treatment. NMIIB (DMSO), 26 cells; NMIIB (Y-27632), 47 cells; NMIIB (RO-3306), 24 cells; active RhoA (DMSO), 30 cells; active RhoA (Y-27632), 55 cells; active RhoA (RO-3306), 35 cells. A ratio higher than 1 means increased intensity after treatments. (C) Measurement of the mitotic cell surface tension by MPA in cells treated with DMSO, 20 μM Y-27632, or 10 μM RO-3306. DMSO, 59 cells; Y-27632, 59 cells; RO-3306, 60 cells. (D) Quantification of the cell height before and after treatment of DMSO, 20 μM Y-27632, or 10 μM RO-3306 with compression. 25 cells in each DMSO or Y-27632 treatment and compression were counted for their cell heights. 27 cells in each DMSO or RO-3306 treatment and compression were counted for their cell heights. Data represent means ± SD. Scale bar, 5 μm.

**Figure S3.**
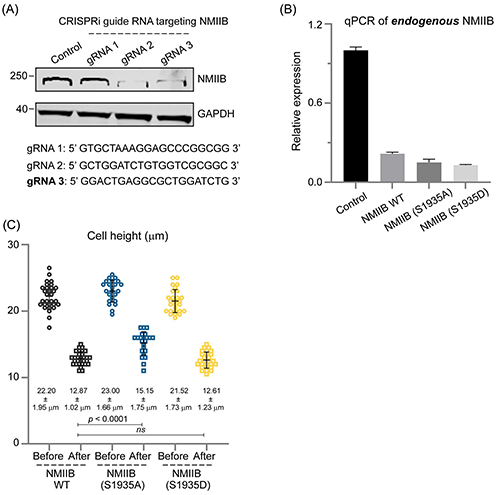
Identification of gRNA sequences that could knockdown NMIIB expression by CRISPRi.(A) Western blotting shows the knockdown efficiency of endogenous NMIIB by CRISPRi using 3 different guiding RNAs (gRNAs). Highlighted gRNA 3 was used in the study to knock down NMIIB. (B) Relative expression of endogenous NMIIB in MCF10A cells, MCF10A cells expressing wild type NMIIB or phosphomutants as determined by qPCR. (C) Quantification of the cell height in cells expressing NMIIB (WT), NMIIB (S1935A), or NMIIB (S1935D) before and after compression. NMIIB (WT), 30 cells; NMIIB (S1935A), 21 cells, NMIIB (S1935D), 26 cells. Data represent means ± SD.

